# Dendritic cells activate pyroptosis and effector-triggered apoptosis to restrict *Legionella* infection

**DOI:** 10.1101/2025.02.13.638189

**Authors:** Víctor R. Vázquez Marrero, Jessica Doerner, Kimberly A. Wodzanowski, Jenna Zhang, Allyson Lu, Frankie D. Boyer, Isabel Vargas, Suzana Hossain, Karly B. Kammann, Madison V. Dresler, Sunny Shin

## Abstract

The innate immune system relies on pattern recognition receptors (PRRs) to detect pathogen-associated molecular patterns (PAMPs) and guard proteins to monitor pathogen disruption of host cell processes. How different immune cell types engage PRR- and guard protein-dependent defenses in response to infection is poorly understood. Here, we show that macrophages and dendritic cells (DCs) respond in distinct ways to bacterial virulence activities. In macrophages, the bacterial pathogen *Legionella pneumophila* deploys its Dot/Icm type IV secretion system (T4SS) to deliver effector proteins that facilitate its robust intracellular replication. In contrast, T4SS activity triggers rapid DC death that potently restricts *Legionella* replication within this cell type. Intriguingly, we found that infected DCs exhibit considerable heterogeneity at the single cell level. Initially, a subset of DCs activate caspase-11 and NLRP3 inflammasome-dependent pyroptosis and release IL-1*β* early during infection. At later timepoints, a separate DC population undergoes apoptosis driven by T4SS effectors that block host protein synthesis, thereby depleting the levels of the pro-survival proteins Mcl-1 and cFLIP. Together, pyroptosis and effector-triggered apoptosis robustly restrict *Legionella* replication in DCs. Collectively, our work suggests a model where Mcl-1 and cFLIP guard host translation in DCs, and that macrophages and DCs distinctly employ innate immune sensors and guard proteins to mount divergent responses to *Legionella* infection.

## Introduction

Upon infection, host cells utilize pattern recognition receptors (PRRs) to recognize pathogen-associated molecular patterns (PAMPs), resulting in immune responses that protect the host (1). However, as many PAMPs are common to both commensal and pathogenic microbes, this mode of recognition alone is insufficient for the immune system to distinguish between avirulent and harmful microbes. To appropriately respond to potentially dangerous microbes, the host additionally senses “patterns of pathogenesis,” such as invasion of pathogens into the host cell cytosol or disruption of cellular processes by microbial effector proteins (2). PAMPs contaminating the host cell cytosol can be detected by a variety of cytosolic innate immune sensors, including those that assemble into multi-protein complexes known as inflammasomes. Meanwhile, virulence factor-mediated disruption of host cell homeostasis is sensed by specific ‘guard’ proteins that normally participate in cellular processes. This latter immune recognition strategy was first described in plants and has been termed “effector-triggered immunity” or “guard immunity” (3, 4). Ultimately, detection of multiple patterns of pathogenesis induces immune responses that can lead to cell death and elimination of the replicative niche for intracellular pathogens. The bacterial virulence activities that induce effector-triggered immunity and how different cell types sense patterns of pathogenesis to mount cell type-specific immune responses against infection remain poorly understood. Using opportunistic bacteria like *Legionella pneumophila* that induce strong protective responses in healthy mammalian hosts can help us learn more about these immune mechanisms, as *Legionella* engages several virulence activities when it infects host cells (5–21).

*Legionella* is naturally found in freshwater environments where it replicates within amoebae (22–27). Humans are infected with *Legionella* upon inhalation of contaminated water droplets, at which point the bacteria can colonize the lungs and cause a severe pneumonia called Legionnaires’ disease (28–30). Although *Legionella* infection can result in significant morbidity and mortality in the elderly or immunocompromised, most healthy individuals clear the bacteria by engaging protective inflammatory pathways (31–34). As *Legionella* has evolved primarily to infect amoebae, this bacterium targets highly conserved eukaryotic processes, but has not adapted to fully evade the mammalian immune system (22–27, 35). Since *Legionella* induces such robust protective responses in mammalian hosts, it serves as an excellent model pathogen to uncover innate immune mechanisms that may otherwise be evaded or disrupted by mammalian-adapted pathogens.

In the lung, the first immune cell type that encounters *Legionella* is the alveolar macrophage, which also serves as the primary replicative niche for the bacteria (30, 36, 37). These cells utilize multiple inflammasomes to detect intracellular *Legionella* infection (38–50). The NAIP5/NLRC4 inflammasome is activated upon recognition of *Legionella* flagellin in the cytosol (38, 39, 42–46, 49, 50), while the NLRP3 inflammasome is activated by T4SS activity (38, 40, 41, 47) and the caspase-11 inflammasome is activated by cytosolic lipopolysaccharide (LPS) (40, 41, 47, 48). Activation of these inflammasomes leads to the processing and activation of caspase-1 and caspase-11, which cleave and activate the pore-forming protein gasdermin D (GSDMD) (51, 52), as well as process and release IL-1 family cytokines (53–58). These events cause an inflammatory and lytic form of cell death known as pyroptosis, which alerts nearby cells to control bacterial infection (59, 60). Additionally, NAIP5/NLRC4-dependent pyroptotic death of infected cells restricts the replicative niche of *Legionella* (39, 42, 45, 46, 61, 62), whereas in unprimed macrophages the NLRP3 and caspase-11 inflammasomes do not limit bacterial replication (40, 41, 47).

Macrophages also activate effector-triggered immunity upon sensing bacterial perturbation of host cell processes. *Legionella* injects over 300 effector proteins via its Dot/Icm type IV secretion system (T4SS) to establish an endoplasmic reticulum (ER)-derived *Legionella*-containing vacuole (LCV) (8, 37, 63–67). These effectors manipulate multiple host cell processes, including membrane trafficking and protein translation (35, 68, 69). *Legionella* has at least 7 effectors (Lgt1-3, SidI, SidL, LegK4, and RavX) that potently block host protein synthesis in macrophages, causing a greater than 95% decrease in host protein synthesis (12, 15, 18, 19, 70–73). Proteins that undergo rapid turnover, like the short-lived nuclear factor-kappa B (NF-κB) negative regulators IκB and A20 are particularly sensitive to generalized disruption of protein synthesis, leading to sustained activation of NF-κB signaling which potently upregulates a subset of inflammatory genes (18). Thus, in this setting, IκB and A20 serve as guards of host protein synthesis. Similarly, another set of pro-survival proteins from the B-cell lymphoma-2 (Bcl-2) family guard host translation during viral infection of human keratinocytes (74). Specifically, simultaneous downregulation of the short-lived anti-apoptotic protein myeloid cell leukemia-1 (Mcl-1) and inactivation of Bcl-XL led to caspase-3-dependent cleavage of gasdermin E (GSDME), which activates pyroptosis to restrict viral replication in human skin cells (74). Whether these pro-survival molecules serve as sensors of translational shutdown in cell types other than keratinocytes or during bacterial infection is unclear.

Many studies have focused on how *Legionella* interacts with macrophages, as they are the primary replicative niche for this bacterium during pulmonary infection (30, 36, 37), but other immune cells in the lung also interact with and respond to *Legionella*. In addition to alveolar macrophages, neutrophils also harbor *Legionella* in mouse models of infection (36). Furthermore, neutrophils and monocytes are required for pro-inflammatory responses and control of pulmonary infection (59, 60, 75–80). At the later stages of infection, a robust T and B cell response is generated (81–89). However, an innate immune cell type that has been largely overlooked is the dendritic cell (DC), which bridges innate and adaptive immune responses in other contexts by acting as antigen presenting cells (90–94). Whether DCs interact with *Legionella* during pulmonary infection remains unclear. In contrast to macrophages, we previously found that DCs are not productively infected during *in vivo* infection (36). Notably, during *in vitro* infection, DCs undergo rapid caspase-3-dependent apoptosis upon detecting *Legionella* T4SS activity, which restricts *Legionella* replication within these cells (95, 96). These findings suggest that in contrast to macrophages, DCs possess unique cell-intrinsic mechanisms that promote apoptosis in response to T4SS-translocated products to restrict bacterial replication.

In this study, we investigated the host pathways and bacterial virulence factors that mediate cell death and restriction of *Legionella* replication in DCs. We found that different populations of infected DCs activate *either* caspase-11 and NLRP3 inflammasome-dependent pyroptosis *or* apoptosis in response to *Legionella* infection. Apoptosis was triggered by *Legionella* T4SS effectors that block host protein synthesis, and *Legionella* lacking these effectors were able to replicate in DCs. We also found that DCs express substantially lower levels of the pro-survival proteins Mcl-1 and cellular FADD-like IL-1β-converting enzyme-inhibitory protein (cFLIP) than macrophages, and effector blockade of host translation led to decreased levels of these proteins. These findings suggest a model where Mcl-1 and cFLIP act as guards of host protein synthesis in DCs, and pathogen interference with host translation leads to decreased levels of these proteins, thus triggering apoptosis. Altogether, we show that DCs activate PRR-triggered pyroptosis and effector-triggered apoptosis to maximally restrict *Legionella* replication. Our findings also highlight major differences in the PRRs and guard proteins used by DCs and macrophages to mount distinct immune responses to *Legionella* infection.

## Results

### Dendritic cells trigger both extrinsic and intrinsic apoptosis in response to *Legionella pneumophila* infection

In contrast to murine bone marrow-derived macrophages (BMDMs), bone marrow-derived dendritic cells (BMDCs) are not permissive for intracellular *Legionella* replication as they undergo rapid mitochondrial apoptosis in response to infection (95, 96). We sought to investigate the bacterial and host factors that mediate this response. To avoid flagellin-mediated activation of the NAIP5/NLRC4 inflammasome, which restricts bacterial replication in both macrophages and DCs and can potentially mask other cell death mechanisms, we employed flagellin-deficient Δ*flaA Legionella* (hereafter T4SS^+^) (38, 39, 42–46, 49, 50, 96). To circumvent any confounding effect of intracellular bacterial replication, we also used thymidine auxotrophic strains of *Legionella* incapable of replicating in the absence of exogenous thymidine. We first assessed cell death kinetics in infected BMDMs and BMDCs by monitoring release of lactate dehydrogenase (LDH) into the extracellular space. As expected, there were significantly higher levels of cell death in BMDCs at 2hr and 4hr post-infection compared to BMDMs, indicating that DCs undergo more robust and rapid cell death than macrophages in response to *Legionella* infection (Fig 1A).

**Figure 1.**
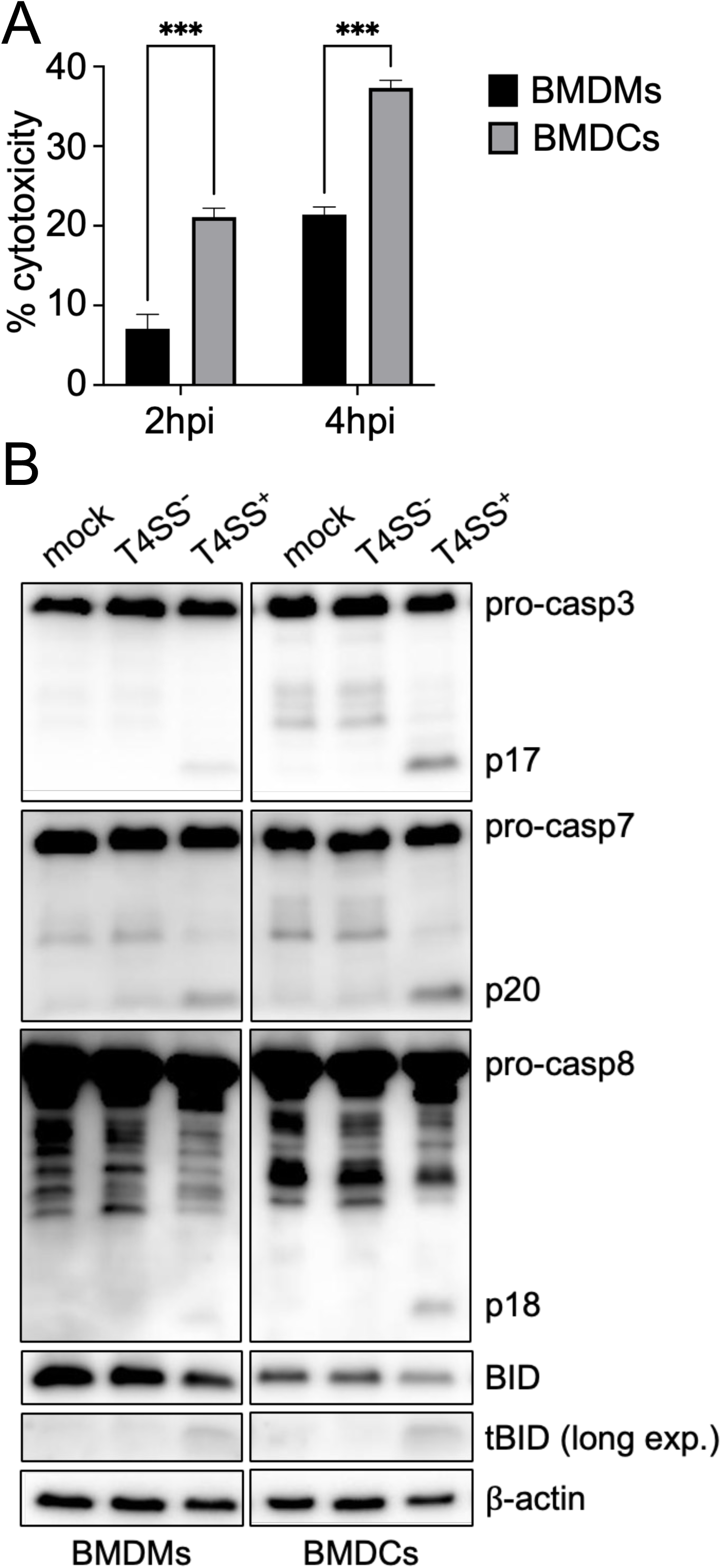
Dendritic cells undergo both cell-extrinsic and intrinsic apoptosis in response to *Legionella* infection. (A) BMDMs and BMDCs were infected with non-replicating Δ*thyA*Δ*flaA Legionella* (T4SS^+^) at a multiplicity of infection (MOI) of 50 for 2 or 4hr. Cytotoxicity was measured by LDH release assay. (B) BMDMs and BMDCs were mock-infected or infected with Δ*thyA*Δ*flaA*Δ*dotA* (T4SS^−^) or T4SS^+^ *Legionella* at an MOI of 50 for 4hr. Immunoblot analysis was performed on cell lysates for caspase-3, caspase-7, caspase-8, and BID with β-actin as a loading control. long exp., long exposure. hpi, hours post-infection. Data shown are representative of at least two (B) or three (A) independent experiments. Graphs show the mean ± SEM of triplicate wells. Data were analyzed by two-way ANOVA with Sidak’s multiple comparisons test; ***, P < 0.001.

We next investigated the apoptotic pathways induced by *Legionella* in DCs. Previous studies found a role for the pro-apoptotic Bcl2-associated X protein (BAX) and Bcl2 antagonist/killer protein (BAK) in activating caspase-3 and restricting *Legionella* replication in BMDCs (96), suggesting that intrinsic mitochondrial pathway of apoptosis is activated in DCs. However, caspase-3 can also be activated by caspase-8 during extrinsic apoptosis. Furthermore, crosstalk between the extrinsic and intrinsic apoptotic pathways can occur through caspase-8-mediated cleavage of BH3-interacting domain death agonist (BID) into its truncated form (tBID), which promotes BAX and BAK activation and mitochondrial permeabilization (97).

Indeed, DCs infected with T4SS^+^ *Legionella* showed increased cleavage of both caspase-8 and BID compared to BMDMs (Fig. 1B). We also observed more robust cleavage of caspase-3 and - 7 in BMDCs compared to BMDMs (Fig. 1B). Importantly, caspase cleavage was dependent on a functional T4SS, as DCs infected with Δ*flaA*Δ*dotA Legionella* lacking the essential T4SS component DotA (hereafter T4SS^−^) did not exhibit cleavage of caspases-3, −7, −8, or BID (Fig 1B). To further address the role of caspase-8, we infected BMDCs deficient for both caspase-8 and receptor-interacting protein kinase 3 (RIPK3), given that caspase-8 single knockout mice exhibit uncontrolled activation of RIPK3-mediated necroptosis and are embryonically lethal (98–101). Immunoblot analysis revealed a partial decrease in caspase-3 and −7 cleavage in *Ripk3^−/−^ Casp8^−/−^*BMDCs infected with T4SS^+^ *Legionella* compared to WT or *Ripk3^−/−^*BMDCs (Fig S1), indicating that caspase-8 is not the sole driver of DC apoptosis and that there is concurrent activation of the intrinsic pathway as previously reported (96). Altogether, these results show that DCs undergo both extrinsic and intrinsic apoptosis in response to *Legionella* T4SS activity.

### *Legionella* T4SS effectors that block host translation trigger apoptosis in dendritic cells

We next aimed to determine the T4SS-dependent bacterial factors responsible for the induction of apoptosis in *Legionella-*infected BMDCs. At homeostasis, apoptosis is inhibited by the pro-survival Bcl-2 family of proteins (102, 103). Some Bcl-2 family members, like Mcl-1, have a short half-life of 0.5 to 3 hours (104, 105). Thus, inhibition of host protein synthesis by chemicals like cycloheximide or microbial toxins leads to rapid depletion of Mcl-1 and other pro-survival proteins, causing some cell types to undergo apoptosis (74, 106–115). Since *Legionella* has at least 7 effectors that potently and redundantly block host translation (12, 15, 18, 19, 70–73), we hypothesized that effector-driven blockade of host protein synthesis triggers apoptosis in DCs.

While *Legionella* inhibits protein translation in macrophages (18), whether it also does so in DCs is unknown. We therefore monitored global protein synthesis in BMDCs with surface sensing of translation (SUnSET) (116), which uses minimal amounts of the aminoacyl tRNA analog puromycin together with anti-puromycin antibodies to detect puromycin incorporation into nascently translated peptides. Immunoblot-based quantification of puromycin incorporation showed that BMDCs infected with T4SS^+^ *Legionella* had reduced levels of protein translation similar to BMDCs treated with cycloheximide, compared to vehicle-treated cells (Fig 2A). In contrast, cells infected with T4SS^−^ *Legionella* unable to inject bacterial effectors had robust levels of protein synthesis. DCs infected with *Legionella* lacking 7 effectors that block host protein synthesis (hereafter T4SS^+^Δ*7*) (72) displayed substantially higher levels of protein synthesis compared to DCs infected with T4SS^+^ *Legionella* (Fig 2A), indicating that *Legionella* uses these effectors to inhibit translation in DCs. The protein synthesis levels of T4SS^+^Δ*7 Legionella*-infected DCs were not fully restored to that of T4SS^−^-infected DCs, in agreement with other studies indicating that T4SS^+^Δ*7 Legionella* still cause a partial block in host protein synthesis (72), suggesting that *Legionella* has additional unknown effectors that block host translation.

**Figure 2.**
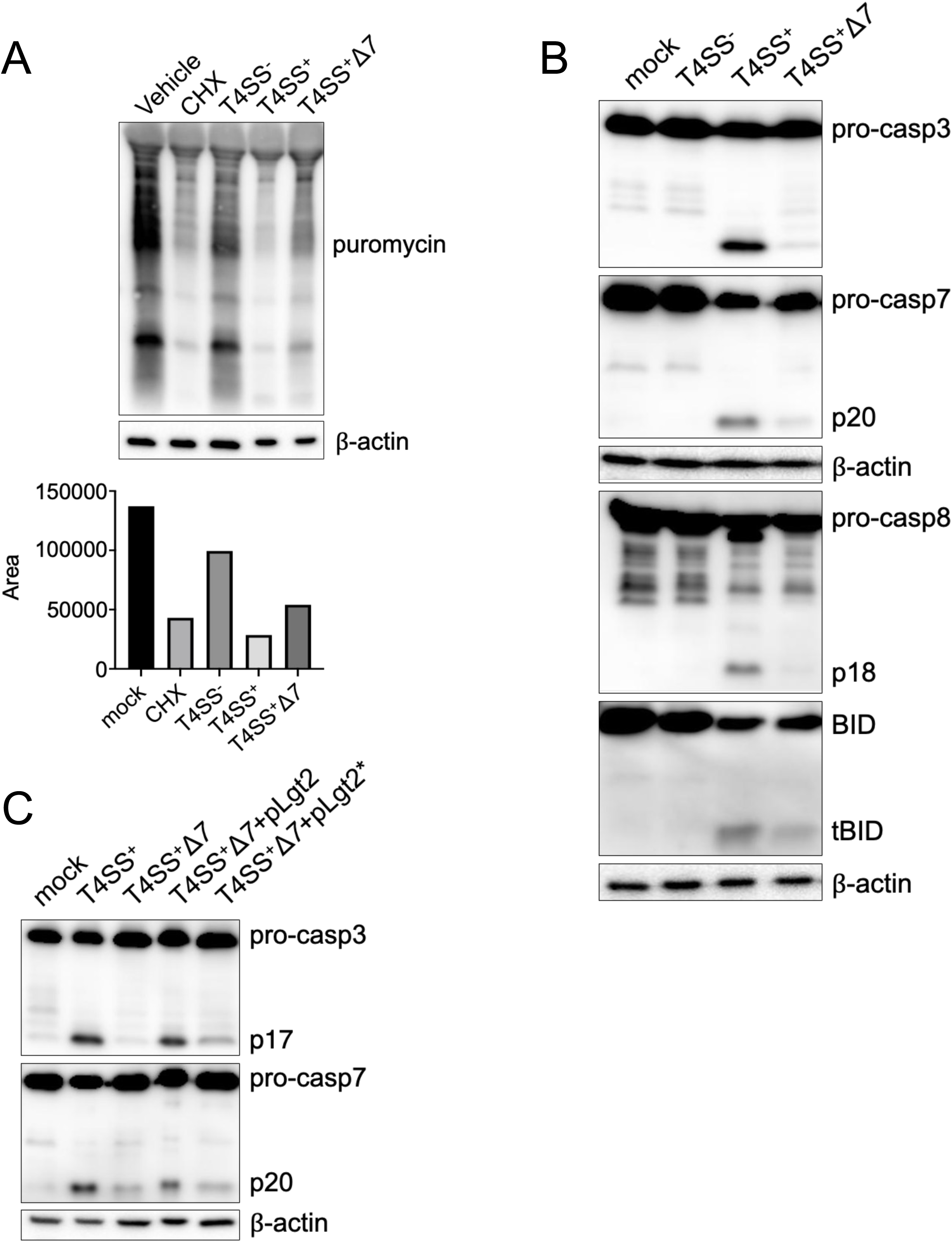
A subset of *Legionella* T4SS effectors block host protein translation and trigger apoptosis in dendritic cells. (A) BMDCs were infected with T4SS^+^ *Legionella* at an MOI of 10 or treated with cycloheximide for 4 hr. 10 ug/mL puromycin was added for 1 hr and cell lysates were harvested. Immunoblot analysis with anti-puromycin antibody was performed on cell lysates. β-actin was used as a loading control. Band density was quantified using ImageJ; bar graph shows corresponding area under the curve. (B) BMDCs were mock-infected or infected with T4SS^−^, T4SS^+^, or Δ*thyA*Δ*flaA*Δ*7 Legionella* (T4SS^+^Δ7) at an MOI of 50 for 4 hr. Immunoblot analysis was performed on cell lysates for caspase-3, caspase-7, caspase-8, BID and β-actin as a loading control. (C) BMDCs were mock-infected or infected with T4SS^+^, T4SS^+^Δ*7*, T4SS^+^Δ*7*+pLgt2, and T4SS^+^Δ*7*+pLgt2* *Legionella*. Lysates were analyzed by immunoblot for caspase-3, caspase-7, and β-actin as a loading control. Data shown are representative of at least three independent experiments.

We next examined whether the *Legionella* effectors that block host protein synthesis are responsible for inducing apoptosis in DCs. DCs infected with T4SS^+^Δ*7 Legionella* had reduced cleavage of caspases-8, −3, −7, and BID compared to cells infected with T4SS^+^ *Legionella* (Fig 2B). Caspase-3 and −7 cleavage was restored by genetically complementing T4SS^+^Δ*7 Legionella* with one of the 7 effectors, *Legionella* glucosyltransferase 2 (Lgt2) (Fig 2C). In contrast, complementing T4SS^+^Δ*7 Legionella* with a catalytically inactive Lgt2 mutant unable to block host protein synthesis (18) did not restore caspase cleavage, indicating that Lgt2’s catalytic activity is required for DCs to activate apoptosis. Altogether, these data show that a subset of *Legionella* T4SS effectors robustly block host protein synthesis and trigger apoptosis in infected DCs.

### Dendritic cells express low levels of anti-apoptotic proteins that are further reduced due to *Legionella* blockade of host protein synthesis

Intrinsic apoptosis is tightly regulated by the balance between pro- and anti-apoptotic proteins, many belonging to the Bcl-2 family of proteins (102, 103). Several Bcl-2 family members, including Mcl-1, have a relatively short half-life, and depletion of these pro-survival factors due to chemical inhibitors or microbial toxins leads to apoptosis in other cell types (74, 106–115). Notably, Mcl-1 and Bcl-XL have been described as guards of host translation during viral infection, as their downregulation or inactivation, respectively, leads to caspase-3 activation and cell death in keratinocytes (74). Extrinsic apoptosis is regulated by an anti-apoptotic caspase-8 homolog called cFLIP. The upregulation of cFLIP in response to NF-κB-activating stimuli and formation of the heterodimeric caspase-8:cFLIP heterodimer blocks apoptosis and promotes cellular survival (117–122). When cFLIP fails to be upregulated due to chemical or bacterial blockade of NF-κB signaling or host translation (123–133), caspase-8 instead forms a homodimer that drives apoptosis (117, 134–136). Given that DCs undergo both intrinsic and extrinsic apoptosis in response to *Legionella* blockade of host protein synthesis, whereas macrophages do not, we hypothesized that DCs basally express lower levels of pro-survival factors than macrophages. Notably, we found that DCs express lower levels of the pro-survival proteins Bcl-XL, Mcl-1, and cFLIP compared to macrophages (Fig. 3A), consistent with the hypothesis that lower expression of pro-survival factors in DCs may render them more susceptible to apoptosis than macrophages in response to bacterial inhibition of protein synthesis.

**Figure 3.**
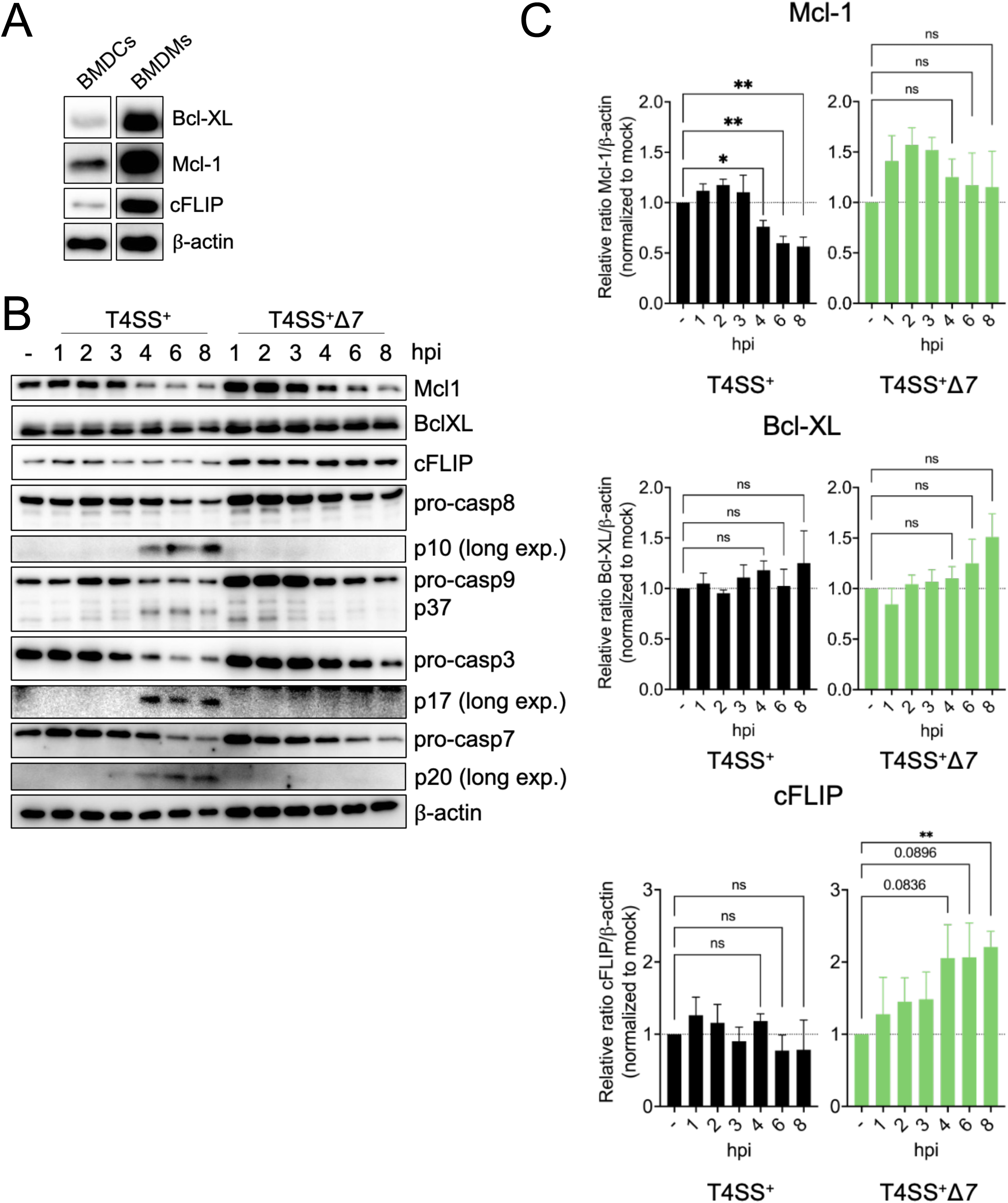
Dendritic cells express low levels of Mcl-1 and cFLIP that are further decreased by *Legionella* T4SS effectors. (A) Immunoblot analysis was performed on the cell lysates of uninfected BMDMs and BMDCs for Bcl-XL, Mcl-1, cFLIP, and β-actin as a loading control. (B) BMDCs were mock-infected (represented as “-“) or infected with T4SS^+^ or T4SS^+^Δ*7 Legionella* at a MOI of 50 for 1, 2, 3, 4, 6, or 8hr post-infection. Immunoblot analysis was performed on cell lysates for Mcl-1, Bcl-XL, cFLIP, caspase-8, caspase-9, caspase-3, caspase-7 and β-actin as a loading control. For cleaved products, lanes from one membrane have been cropped and moved to depict the appropriate conditions. No changes were made to the original image during the editing. (C) Three independent experiments from B were quantified using ImageJ for Bcl-XL, Mcl-1, and cFLIP and the average was graphed for each protein. long exp., long exposure. hpi, hours post-infection. Graphs show the mean ± SEM of triplicate wells. Data shown are representative of at least (A) two and (B) three independent experiments. Data in C were analyzed by unpaired t test; **, P < 0.01; *, P < 0.05; ns, not significant.

We next determined whether the levels of these pro-survival factors are impacted by *Legionella* effector-mediated blockade of host translation. DCs infected with T4SS^+^ *Legionella* had Mcl-1 levels that were significantly decreased by 4 hours post-infection compared to uninfected cells (Fig. 3B and 3C). In contrast, in DCs infected with T4SS^+^Δ*7 Legionella,* although Mcl-1 levels were slightly decreased, they remained at higher levels (Fig. 3B and 3C). Bcl-XL levels were unaffected by the *Legionella-*induced block of host protein synthesis (Fig. 3B and 3C), potentially due to its long half-life of 20 hours (137). cFLIP levels in DCs infected with T4SS^+^ *Legionella* were similar to uninfected DCs, whereas cFLIP levels were significantly upregulated in T4SS^+^Δ*7 Legionella*-infected cells compared to T4SS^+^ *Legionella-*infected or uninfected cells, suggesting that the effector-mediated block of host translation prevents cFLIP upregulation. In T4SS^+^ *Legionella-*infected DCs, cleavage of the apoptotic caspases-8, −9, −3, and −7 began at 4 hours post-infection, coinciding with the decrease in Mcl-1 levels (Fig. 3B and 3C). In contrast, there was no apoptotic caspase cleavage in T4SS^+^Δ*7 Legionella*-infected DCs. Taken together, these data indicate that in contrast to macrophages, DCs express lower levels of the critical pro-survival factors Mcl-1 and cFLIP that regulate intrinsic and extrinsic apoptosis, potentially explaining why DCs undergo rapid apoptosis in response to *Legionella* whereas macrophages do not. Additionally, these results suggest a model where pro-survival factors are rapidly depleted or fail to be upregulated in DCs due to the *Legionella* effector-mediated block of host translation, thereby directing DCs to die by apoptosis.

### Dendritic cells undergo caspase-1-, caspase-11-, and gasdermin D-dependent pyroptosis early during *Legionella* infection

Our analysis revealed that the *Legionella* effectors that block host protein synthesis trigger apoptotic caspase cleavage at around 4 hours post-infection (Fig 3B). However, we observed cell death as early as 2 hours post-infection (Fig 1A), suggesting that other cell death responses independent of these effectors are occurring in DCs. To more rigorously assess cell death kinetics, we measured propidium iodide (PI) uptake over time in DCs infected with T4SS^+^ or T4SS^+^Δ*7 Legionella*. We observed only a partial decrease in cell death in T4SS^+^Δ*7*-infected DCs compared to T4SS^+^-infected DCs (Fig S2), despite the absence of apoptotic caspase cleavage in these cells (Fig 3B). These data indicate that effector-triggered apoptosis does not account for all of the cell death occurring in infected DCs.

We next investigated what additional forms of cell death occur in *Legionella-*infected DCs. In macrophages, *Legionella* activates the NAIP5/NLRC4 and caspase-11 inflammasomes (38–50), which lead to caspase-1 and GSDMD activation, IL-1 cytokine release, and pyroptosis (138). In DCs, *Legionella* also activate the NAIP5/NLRC4 inflammasome (96), but whether additional inflammasomes are activated is unknown. In this study, we use flagellin-deficient *Legionella,* eliminating the contribution of the NAIP5/NLRC4 inflammasome. Thus, we hypothesized that flagellin-deficient *Legionella* trigger caspase-1 and caspase-11-dependent pyroptosis in DCs. To test this, we infected BMDCs from WT, *Casp11^−/−^,* or *Casp1^−/−^Casp11^−/−^* mice. Cell death within the first 4 hours of infection mainly depended on caspase-11, as measured by PI uptake and LDH release (Fig 4A and 4B), with a modest additional decrease in cell death in *Casp1^−/−^Casp11^−/−^* BMDCs compared to *Casp11^−/−^* BMDCs. DCs infected with T4SS^+^ *Legionella* had low levels of IL-1 production due to the block in host protein synthesis (Fig S3) (5, 139, 140). Therefore, we infected BMDCs with T4SS^+^Δ*7 Legionella* to assess the roles of caspases-1 and −11 in IL-1α and IL-1β release. Deletion of caspase-11 alone resulted in a significant decrease in IL-1α and IL-1β release relative to wildtype, while deletion of both caspases 1 and 11 fully abrogated IL-1 levels (Fig 4C). These data indicate that caspase-1 and-11 work together to promote optimal IL-1 release from DCs during *Legionella* infection.

**Figure 4.**
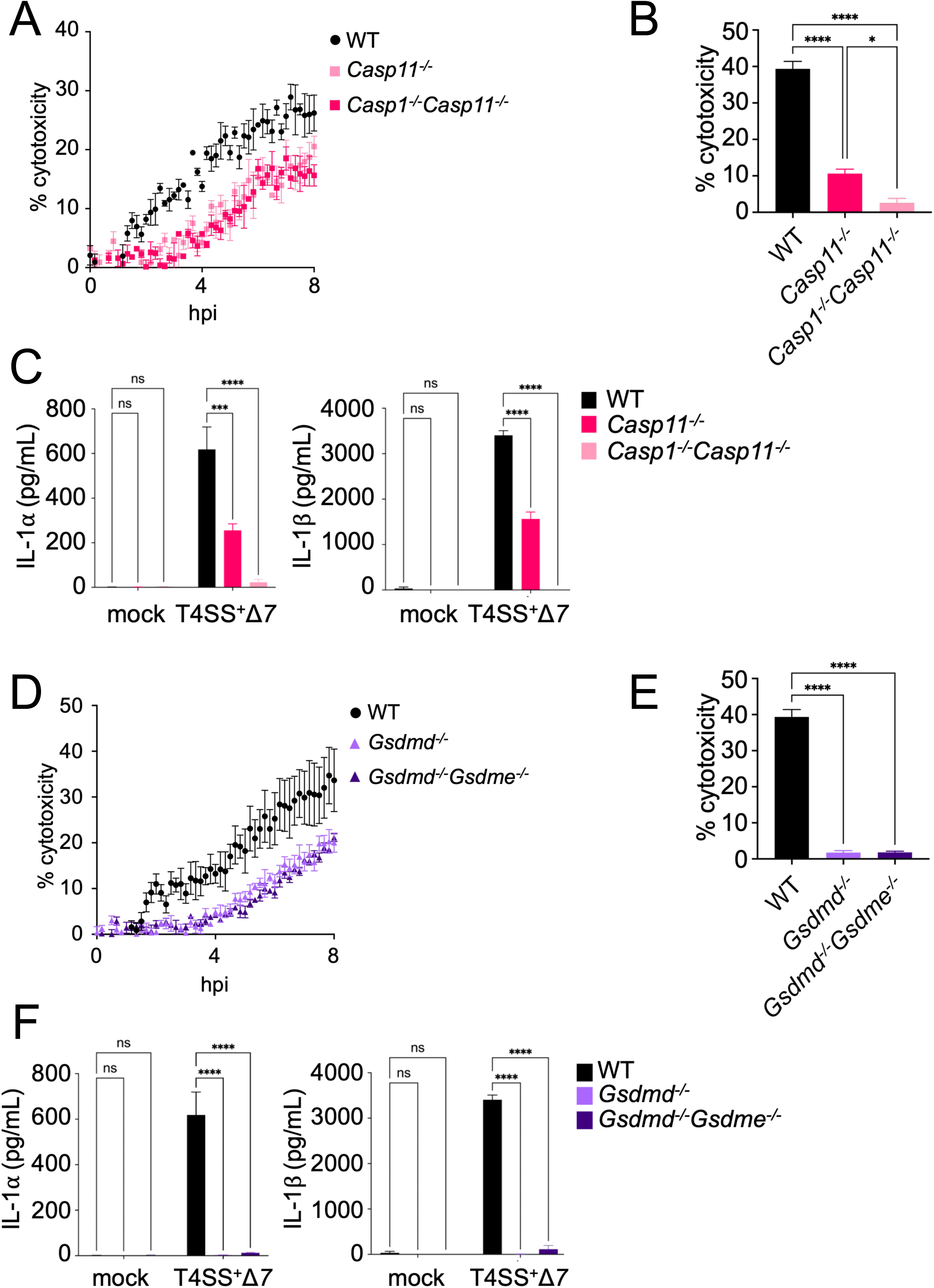
Dendritic cells undergo rapid pyroptosis in response to *Legionella* infection. (A-C) WT, *Casp11^−/−^*, or *Casp1^−/−^Casp11^−/−^* BMDCs were mock-infected (C) or infected with T4SS^+^ (A, B) or T4SS^+^Δ*7 Legionella* (C) at an MOI of 50. Cytotoxicity was measured by PI uptake assay (A) or LDH release assay at 4hr (B). Cytokine release was measured at 4hr using ELISA (C). (D-F) WT, *Gsdmd^−/−^*, or *Gsdmd^−/−^Gsdme^−/−^*BMDCs were mock-infected (F) or infected with T4SS^+^ (D, E) or T4SS^+^Δ*7 Legionella* (F) at an MOI of 50. Cytotoxicity was measured by PI uptake assay (D) or LDH release assay at 4hr (E). Cytokine release was measured at 4hr using ELISA (F). hpi, hours post-infection. Data shown are representative of at least three independent experiments. Graphs show the mean ± SEM of triplicate wells. Data were analyzed by two-way ANOVA with Sidak’s multiple comparisons test; ****, P < 0.0001; ***, P < 0.001; *, P <0.05; ns, not significant.

We next interrogated the role of terminal gasdermins in mediating DC pyroptosis during *Legionella* infection. Caspase-1 and caspase-11 cleave the cytosolic protein GSDMD, liberating its active pore-forming N-terminal domain, which then oligomerizes and forms a pyroptotic pore from which the cell can release IL-1 family cytokines and damage-associated molecular patterns (51–58, 141, 142). To assess the role of GSDMD in DC pyroptosis, we infected *Gsdmd*^−/−^ BMDCs with T4SS^+^ *Legionella*. We observed substantially less cell death, as measured by PI uptake and LDH release, in *Gsdmd*^−/−^ BMDCs within the first 4hr of infection compared to WT BMDCs (Fig 4D and 4E). However, there were still substantial levels of cell death remaining in infected *Gsdmd^−/−^* BMDCs (Fig 4D). Given that in other circumstances, GSDME can mediate pyroptosis and compensate for GSDMD deficiency (47, 143–148), we additionally infected *Gsdmd^−/−^Gsdme*^−/−^ BMDCs. However, we did not see a further reduction in cell death (Fig 4D and 4E). There was also nearly complete abrogation of IL-1α and IL-1β release in infected *Gsdmd^−/−^* and *Gsdmd^−/−^Gsdme*^−/−^ BMDCs (Fig 4F), indicating that GSDMD, not GSDME, is the primary driver of pyroptosis in *Legionella*-infected DCs. Collectively, these data indicate that in addition to T4SS effector-triggered apoptosis, DCs activate caspase-11- and GSDMD-dependent pyroptosis early during *Legionella* infection.

Tumor necrosis factor (TNF) signaling can contribute to both caspase-11-dependent pyroptosis and caspase-8-mediated apoptosis (47, 136, 149, 150). We therefore determined the role of TNF in promoting cell death in *Legionella*-infected DCs. Notably, cell death levels were significantly reduced in *Tnf^−/−^* BMDCs infected with T4SS^+^ *Legionella* compared to infected WT BMDCs (Fig S4A and S4B). We also observed delayed GSDMD processing into its active p32 fragment in infected *Tnf^−/−^* BMDCs, but found no defect in apoptotic caspase cleavage (Fig S4C and S4D), indicating that TNF facilitates pyroptosis but does not contribute to apoptosis in *Legionella*-infected DCs. We previously demonstrated that TNF licenses macrophages to upregulate and rapidly activate the caspase-11 inflammasome in response to *Legionella* infection (47). However, we observed similar levels of caspase-11 expression in WT and *Tnf^−/−^* BMDCs infected with T4SS^+^ *Legionella* (Fig S4C), suggesting that TNF promotes pyroptosis through other mechanisms in DCs.

### Dendritic cells activate the NLRP3 inflammasome to mediate IL-1*β* release during *Legionella* infection

Our data indicated that although caspase-1 does not play a major role in cell death in *Legionella-*infected DCs, it is required for IL-1β release (Fig 4C). We therefore sought to investigate the pathways activating caspase-1. Previous studies have shown that caspase-11-dependent GSDMD pore formation in macrophages leads to potassium efflux, which triggers the formation of the non-canonical NLRP3 inflammasome containing ASC and pro-caspase-1 (40, 151–155). Pro-caspase-1 is then processed into its mature form, which promotes the cleavage and release of IL-1*β*. Given that *Legionella* activates both the canonical and non-canonical NLRP3 inflammasomes in mouse macrophages (38, 40, 41, 47) and our data indicate that both caspases-1 and −11 are required by *Legionella-*infected DCs for IL-1*β* release, we hypothesized that the NLRP3 inflammasome mediates caspase-1-dependent IL-1 cytokine release in infected DCs downstream of caspase-11 activation. Immunoblot analysis showed that caspase-1 cleavage was substantially decreased in T4SS^+^ *Legionella*-infected *Nlrp3^−/−^* BMDCs, indicating that the NLRP3 inflammasome is required for caspase-1 activation (Fig 5A). Furthermore, we observed nearly complete abrogation of IL-1*β* release in T4SS^+^Δ*7 Legionella*-infected *Nlrp3^−/−^*or *Casp1^−/−^* BMDCs compared to WT, indicating that the NLRP3 inflammasome is the primary driver of IL-1*β* release (Fig 5B). However, there was only a partial decrease in IL-1*α* release in *Nlrp3^−/−^* or *Casp1^−/−^* BMDCs. Given that IL-1*α* release is substantially decreased in BMDCs deficient for both caspase-1 and caspase-11 (Fig 4C), these data suggest that caspase-1 and caspase-11 individually contribute to IL-1*α* release. We did not observe any defects in cell death in *Nlrp3^−/−^* or *Casp1^−/−^* BMDCs infected with T4SS^+^ *Legionella* (Fig 5C), indicating that while NLRP3 and caspase-1 primarily drive IL-1*β* release, they are not required for DC death during *Legionella* infection.

**Figure 5.**
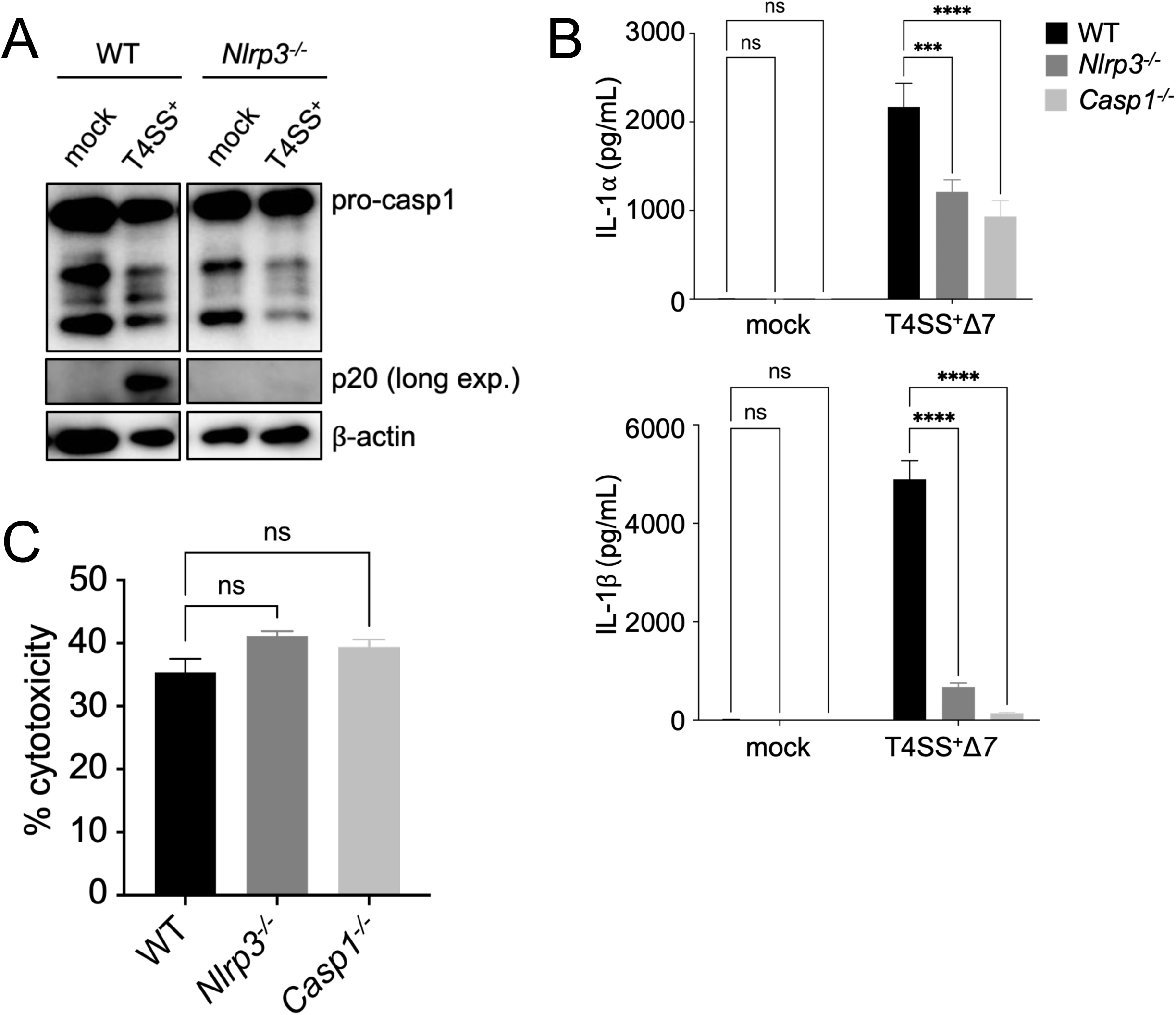
Dendritic cells activate the NLRP3 inflammasome to promote caspase-1 activation and IL-1 release. (A) WT and *Nlrp3^−/−^* BMDCs were mock-infected or infected with T4SS^+^ *Legionella* at a MOI of 50 for 4hr. Immunoblot analysis was performed on cell lysates for caspase-1 and β-actin as a loading control. Lanes from one membrane have been cropped and moved to depict the appropriate conditions. No changes were made to the original image during the editing. (B-C) WT, *Nlrp3^−/−^*, and *Casp1^−/−^* BMDCs were mock-infected (B) or infected with T4SS^+^Δ*7* (B) or T4SS^+^ (C) *Legionella* at an MOI of 50 for 4hr. Cytokine release was measured using ELISA (B). Cytotoxicity was measured by LDH release assay (C). long exp., long exposure. hpi, hours post-infection. Data shown are representative of at least three independent experiments. Graphs show the mean ± SEM of triplicate wells. Data were analyzed by two-way ANOVA with Sidak’s multiple comparisons test; ****, P < 0.0001; ***, P < 0.001. ns, not significant.

### Dendritic cells activate either pyroptosis or effector-triggered apoptosis to restrict *Legionella* infection

We found that DCs activate both pyroptosis and effector-triggered apoptosis in response to *Legionella* infection, yet the relative kinetics of these pathways remained unclear. We therefore assessed the relative timing of pyroptosis and apoptosis in WT DCs infected with T4SS^+^ or T4SS^+^Δ*7 Legionella*. We observed rapid induction of pyroptosis within 1 hour of T4SS^+^ or T4SS^+^Δ*7 Legionella* infection, as evidenced by GSDMD processing into its active p32 fragment (Fig 6A). Later, at 3 to 4 hours post-infection, we observed caspase-7 cleavage into the p20 form following infection with T4SS^+^ *Legionella,* but not T4SS^+^Δ*7 Legionella*, indicating that apoptosis is triggered following pyroptosis during T4SS^+^ *Legionella* infection. At 4 hours following T4SS^+^ *Legionella* infection, we also observed GSDMD processing into its inactive p23 and p43 fragments, possibly due to activity of caspases-3 and −7 (156, 157). In contrast, we observed more GSDMD p32 cleavage and the absence of p23 and p43 fragments in DCs infected with T4SS^+^Δ*7 Legionella,* suggesting that caspases-3 and −7 limit formation of the active GSDMD p32 form during infection with T4SS^+^ *Legionella*.

**Figure 6.**
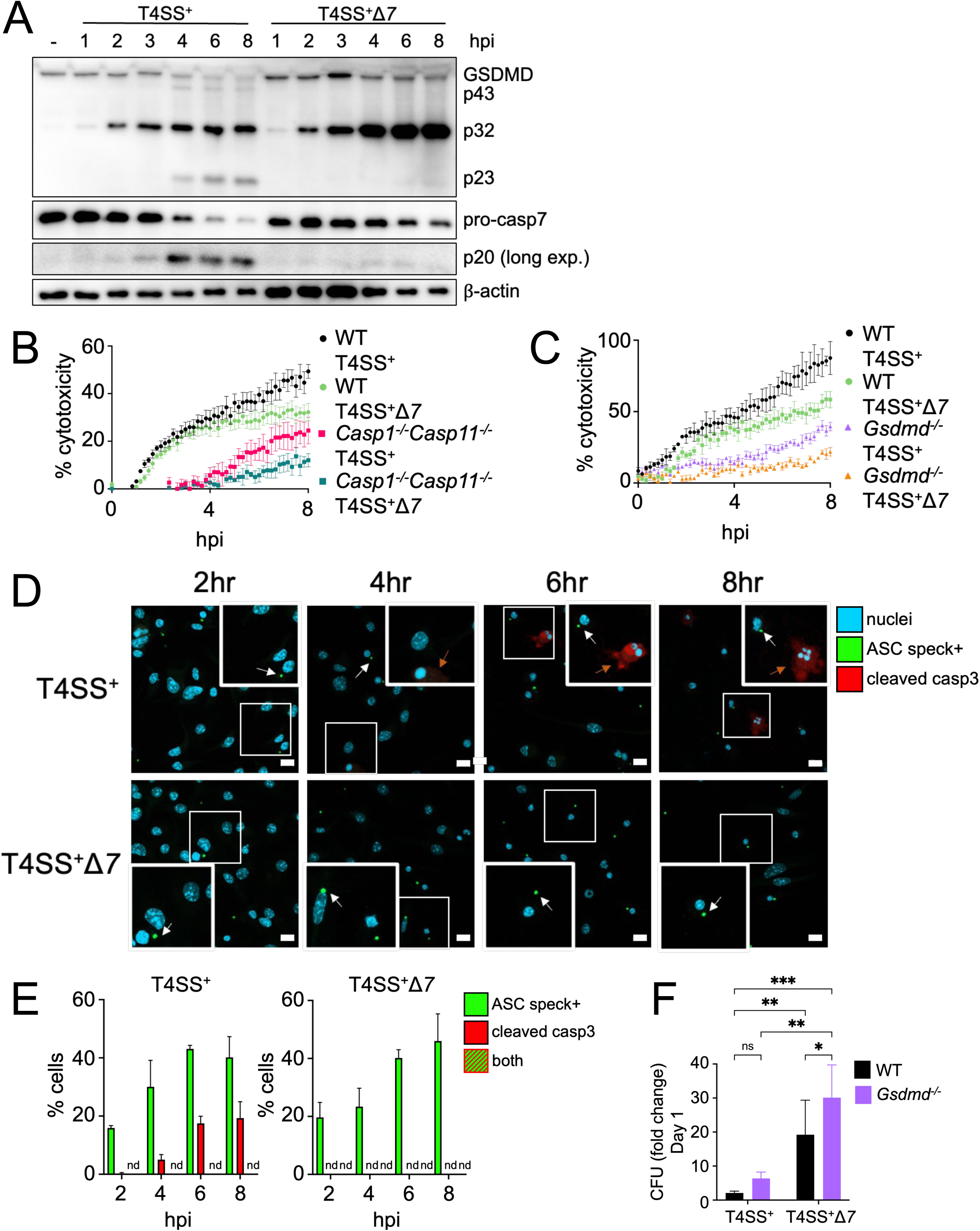
Dendritic cells activate either pyroptosis or effector-triggered apoptosis to restrict *Legionella* infection. (A) BMDCs were mock-infected (represented as “-“) or infected with T4SS^+^ or T4SS^+^Δ*7 Legionella* at an MOI of 50 for 1, 2, 3, 4, 6, or 8 hr post-infection. Immunoblot analysis was performed on cell lysates for GSDMD, caspase-7 and β-actin as a loading control. (B) WT or *Casp1^−/−^Casp11^−/−^* BMDCs were infected with T4SS^+^ or T4SS^+^Δ*7 Legionella* at an MOI of 50. Cytotoxicity was measured by PI uptake assay. (C) WT or *Gsdmd^−/−^* BMDCs were infected with T4SS^+^ or T4SS^+^Δ*7 Legionella* at an MOI of 50. Cytotoxicity was measured by PI uptake assay. (D) ASC-citrine reporter BMDCs were infected with T4SS^+^ or T4SS^+^Δ*7 Legionella* at an MOI of 50 for 2, 4, 6, or 8 hr post-infection and prepared for confocal microscopy. Cells were stained for cleaved caspase-3 (red) and nuclei (blue) with Hoechst. White arrows point ASC speck-positive cells and orange arrows point cleaved caspase-3-positive cells. Insets represent a 1.75x zoom. Scale bars =10 µm. (E) ASC speck-positive (green) and cleaved caspase-3-positive cells from (D) were quantified by counting at least 100 total cells per condition for each independent experiment. (F) WT and *Gsdmd^−/−^* BMDCs were infected with T4SS^+^ or T4SS^+^Δ*7 Legionella* at an MOI of 10. The fold change in colony forming units (CFU) was quantified at 1 day post-infection. long exp., long exposure. hpi, hours post-infection. nd, not detected. (A, B, C, E) Data shown are representative of at least three independent experiments. (D, F) Data represent the pooled results of at least 3 independent experiments. Graphs show the mean ± SEM of triplicate wells. Data were analyzed by two-way ANOVA with Sidak’s multiple comparisons test; **, P < 0.01; *, P < 0.05; ns, not significant.

As a complementary approach, we further assessed the kinetics of pyroptosis and apoptosis in infected DCs by measuring PI uptake in WT or *Casp1/11^−/−^* DCs infected with either T4SS^+^ or T4SS^+^Δ*7 Legionella*. *Casp1/11^−/−^* DCs did not undergo cell death within the first 4 hours of T4SS^+^ *Legionella* infection, but underwent cell death at later timepoints, whereas WT DCs exhibited rapid cell death within one hour post-infection (Fig 6B). WT BMDCs infected with T4SS^+^Δ*7 Legionella* also underwent rapid cell death at levels comparable to WT DCs infected with T4SS^+^ *Legionella* within the first 4 hours of infection, but had reduced levels of cell death at later timepoints. In contrast, *Casp1/11^−/−^* DCs infected with T4SS^+^Δ*7 Legionella* showed minimal cell death at both early and late timepoints.

To further define the kinetics of GSDMD-dependent pyroptosis and effector-triggered apoptosis, we next infected WT or *Gsdmd^−/−^* DCs with T4SS^+^ or T4SS^+^Δ*7 Legionella*. *Gsdmd^−/−^*DCs infected with T4SS^+^ *Legionella* had a nearly complete reduction in cell death at early timepoints and then underwent some cell death after 4 hours post-infection compared to WT DCs infected with T4SS^+^ *Legionella* (Fig 6C). Consistent with our previous observations in *Casp1/11^−/−^* DCs, we observed a nearly complete abrogation of cell death in T4SS^+^Δ*7 Legionella*-infected *Gsdmd^−/−^* DCs compared to the other conditions. Altogether, these data indicate that DCs activate pyroptosis first followed by effector-triggered apoptosis, and that both of these pathways account for the majority of the programmed death occurring in DCs.

Given that the kinetics of pyroptosis and apoptosis partially overlap in T4SS^+^ *Legionella-* infected DCs, we considered whether individual DCs simultaneously experience both pyroptosis and apoptosis or whether pyroptosis and apoptosis occur in distinct subpopulations of cells during infection. During pyroptosis, NLRP3, ASC, and caspase-1 form an inflammasome complex that can be visualized as a large perinuclear speck by microscopy (158–164). We therefore imaged ASC specks using BMDCs derived from mice expressing an ASC-citrine fusion protein, whereas apoptotic cells were identified with antibody-based staining for cleaved caspase-3. Confocal microscopy analysis revealed that DCs infected with T4SS^+^ or T4SS^+^Δ*7 Legionella* contained ASC specks starting at 2 hours post infection, and the percentage of cells positive for ASC specks steadily increased over 8 hours post-infection (Fig. 6D and 6E). Cells positive for cleaved caspase-3 began appearing at 4 hours after infection with T4SS^+^ *Legionella,* but not during infection with T4SS^+^Δ*7 Legionella* (Fig. 6D and 6E), indicating that the presence of cleaved caspase-3 is dependent on the effectors that block host protein synthesis. Notably, we did not find individual T4SS^+^ *Legionella*-infected DCs positive for both ASC specks and cleaved caspase-3 (Fig 6D and Fig 6E). These data indicate that infected DCs do not simultaneously undergo pyroptosis and apoptosis, but rather they exhibit heterogeneity at the single cell level, even when there is apparent kinetic overlap between the two pathways. Thus, a subset of DCs initially engage rapid pyroptosis, whereas another subset of DCs eventually undergo apoptosis in response to the T4SS effectors that block host translation.

We next asked whether both pyroptosis and effector-triggered apoptosis contribute to the restriction of *Legionella* replication within DCs by infecting WT or *Gsdmd^−/−^* BMDCs with T4SS^+^ or T4SS^+^Δ*7 Legionella*. Although *Legionella* replication within *Gsdmd^−/−^* DCs infected with T4SS^+^ *Legionella* was slightly higher than WT DCs infected with T4SS^+^ *Legionella*, it was not statistically significant (Fig 6F). In contrast, WT DCs infected with T4SS^+^Δ*7 Legionella* exhibited a significant increase in bacterial replication compared to WT DCs infected with T4SS^+^ *Legionella* (Fig 6F). We observed the greatest fold change in bacterial replication within *Gsdmd^−/−^* DCs infected with T4SS^+^ Δ*7 Legionella*, which do not activate pyroptosis nor effector-triggered apoptosis, and this increase was significantly higher compared to all other conditions (Fig 6F). Collectively, these results indicate that effector-triggered apoptosis restricts *Legionella* replication to a greater extent than pyroptosis, but that both are required for maximal restriction of *Legionella* by DCs.

## Discussion

In this study, we employed *Legionella pneumophila* to understand how DCs undergo cell death to restrict intracellular bacterial replication. Our findings show that DCs engage extrinsic and intrinsic apoptosis as an effector-triggered immune response to *Legionella* infection. DC apoptosis was triggered by a subset of *Legionella* T4SS effectors that block host protein synthesis, leading to decreased levels of the pro-survival proteins Mcl-1 and cFLIP. We also found that DCs express lower baseline levels of the pro-survival factors Mcl-1, Bcl-XL, and cFLIP compared to macrophages, which may account for why DCs undergo rapid apoptosis in response to the host translational block whereas macrophages do not. In addition to apoptosis, infected DCs underwent rapid caspase-11 and NLRP3 inflammasome activation in response to *Legionella*. There was considerable heterogeneity at the single-cell level, as individual DCs activated either pyroptosis or apoptosis. Finally, both pyroptosis and effector-triggered apoptosis were required for maximal restriction of *Legionella* replication in DCs (Fig 7). Thus, DCs employ both effector-triggered apoptosis and pyroptosis to control *Legionella* infection.

**Figure 7.**
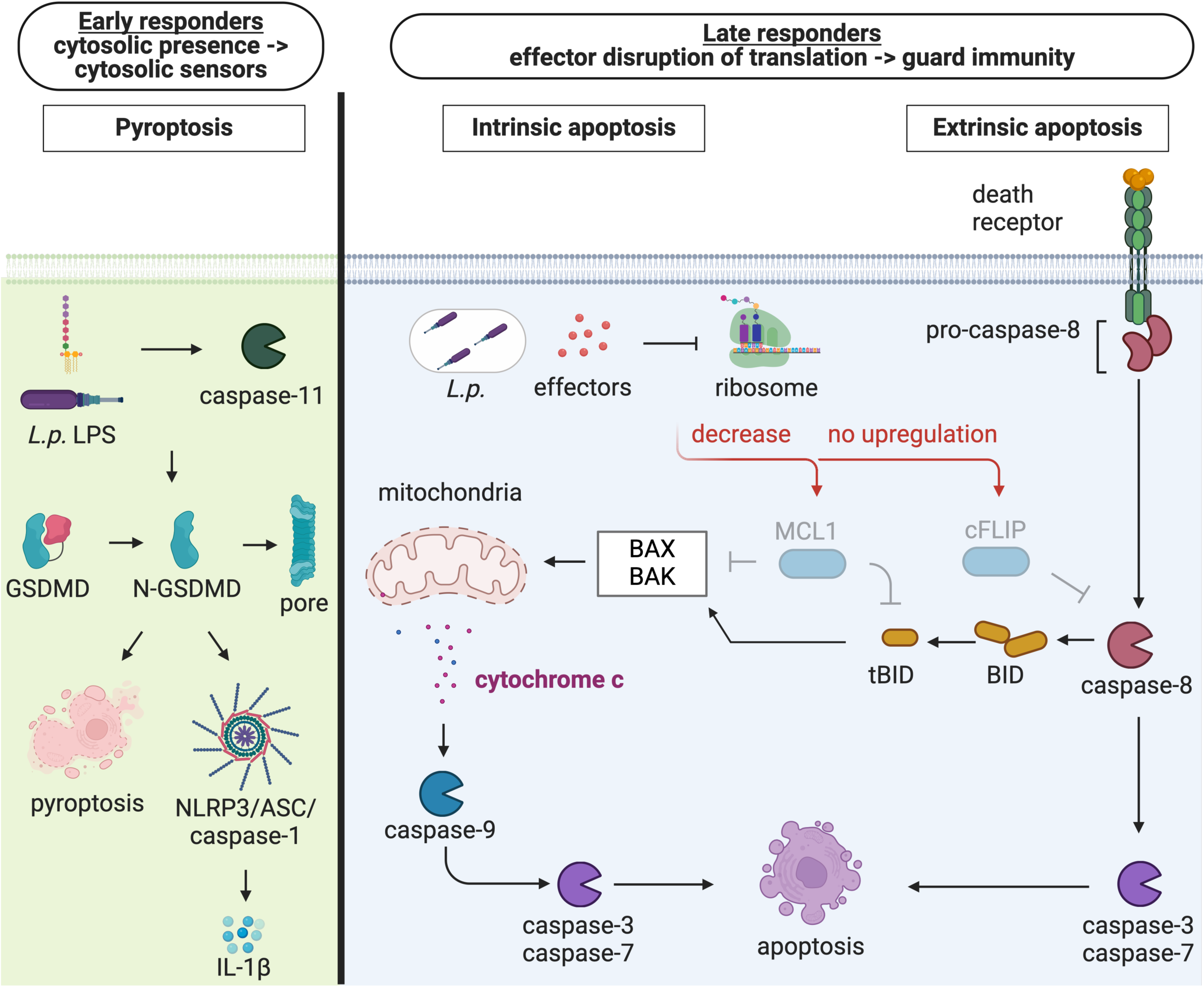
Model of dendritic cell activation of pyroptosis and effector-triggered apoptosis in response to *Legionella* infection. At the early stages of infection (left), a subset of DCs activate caspase-11-dependent pyroptosis in response to *Legionella* LPS in the host cytosol. At the later stages of infection (right), in another subset of DCs, *Legionella* translocates T4SS effectors that block host protein synthesis, which lead to decreased levels of pro-survival molecules that guard host translation, resulting in apoptosis. Figure created in https://BioRender.com

The ‘guard’ hypothesis originally proposed in plant immunity states that host proteins monitor and sense host cellular processes that are common targets of pathogen effectors, thereby acting as guards of homeostasis (2–4, 165, 166). One such cellular process that our study and others have shown to be monitored is host protein synthesis (18, 74, 139, 140, 167–171). Pro-survival proteins of the Bcl-2 family and cFLIP have been previously described to guard host translation against chemical or pathogen inhibitors (104, 105, 109–115, 172, 173). Mcl-1 and cFLIP have short half-lives of 30 minutes and 2 hours, respectively, and thus their protein levels are rapidly depleted upon translational inhibition by cycloheximide, leading to apoptosis (104, 133). In keratinocytes, viral inhibition of translation depletes Mcl-1 and inactivates Bcl-XL, triggering caspase-3- and gasdermin E-mediated pyroptosis that promotes control of viral infection (74). Furthermore, cFLIP expression is induced by NF-κB-activating stimuli, and chemical inhibitors of NF-κB signaling or protein synthesis prevent cFLIP upregulation, thereby triggering apoptosis (123–133). We found that *Legionella* inhibition of host translation led to decreased Mcl-1 and cFLIP levels that coincided with the induction of apoptosis in DCs. Our findings suggest that Mcl-1 and cFLIP act as guards of host protein synthesis and sense bacterial inhibition of host translation in DCs. It will be of interest to uncover the role of these translational guard proteins during infection with other bacterial pathogens in DCs and other cell types.

Our study indicates that DCs possess unique properties that enable them to activate pyroptosis and apoptosis more quickly than macrophages to bacterial infection. Unprimed macrophages normally do not activate the caspase-11 inflammasome until several hours following *Legionella* infection (40, 41, 47). In contrast, we found that DCs undergo rapid caspase-11-dependent pyroptosis as early as 1-hour post-infection. Interestingly, this rapid response was dependent on TNF signaling, similar to LPS- or TNF-primed macrophages which upregulate caspase-11 and other host factors that promote rapid caspase-11 inflammasome activation within 1-hour post-infection (40, 41, 47). However, we did not observe differences in caspase-11 expression in *Legionella*-infected DCs lacking TNF, suggesting that TNF promotes rapid pyroptosis through other mechanisms. Moreover, we found that infected DCs also undergo rapid apoptosis in response to effector blockade of protein synthesis, whereas *Legionella-* infected macrophages do not. Instead, previous studies have shown that in *Legionella-*infected macrophages, effector blockade of host translation leads to increased NF-κB signaling and inflammatory gene expression (18). We show that DCs express lower levels of the pro-survival molecules Bcl-XL, Mcl-1, and cFLIP than macrophages, suggesting a mechanism by which DCs are more sensitive than macrophages to apoptosis in response to perturbations in protein synthesis. Work by Speir *et al.* has shown that macrophages degrade Mcl-1 at late timepoints during *Legionella* infection, but that Bcl-XL remains stable and at high levels (174). Speir *et al*. propose that these high Bcl-XL levels prevent macrophage apoptosis, thus allowing *Legionella* replication. In support, depletion or pharmacological inhibition of Bcl-XL results in macrophage apoptosis and restriction of *Legionella* replication (174). Taken together, our work suggests that DCs are uniquely poised to undergo apoptosis in response to blockade of protein synthesis due to their specific balance of pro- and anti-apoptotic proteins. It will be of interest to undertake future studies aimed at examining whether infected DCs undergo pyroptosis or effector-triggered apoptosis infection during *in vivo Legionella* infection.

It is unclear why DCs and macrophages behave so disparately upon intracellular bacterial infection. As antigen-presenting cells, DCs traffic from the site of infection to the lymph nodes, where they activate adaptive immunity (90–94, 175–179). If DCs harbor viable pathogens during infection, they may act as a Trojan horse and transport and spread pathogens to other sites which would be disadvantageous for the host. Perhaps DCs have evolved to rapidly die in response to bacterial pathogens in order to prevent bacterial dissemination. This finding may extend to other pathogens, as *Salmonella* Typhimurium, *Mycobacterium tuberculosis*, *Listeria monocytogenes*, vaccinia virus, and influenza virus are reported to infect but not replicate within DCs (180–184). Thus, DCs may represent a cellular niche designated to inhibit pathogen replication and dissemination during infection. In turn, some pathogens may have evolved strategies to replicate within DCs and promote their dissemination.

Our data indicate that individual *Legionella-*infected DCs undergo either apoptosis or pyroptosis, but not both (Fig 6C). It is unclear what determines whether a given DC will die by one pathway or another. Perhaps it is due to differential expression of anti-apoptotic factors or inflammasome components at the single-cell level. Alternatively, GM-CSF-derived bone marrow cultures are heterogeneous, as they contain BMDCs in several differentiation stages as well as monocyte-derived macrophages (185, 186). Thus, it is possible that depending on their cellular differentiation state, an individual cell may die by either pyroptosis or apoptosis. Alternatively, perhaps differences in the rates of effector translocation by individual bacteria lead to the triggering of apoptosis or pyroptosis in a given infected DC. Additional studies are needed to better understand the mechanisms underlying the heterogeneity of cell death decisions in *Legionella-*infected BMDCs at a single cell level. It would also be of interested to investigate how different DC subsets interact with *Legionella* during *in vitro* and *in vivo* infection.

One of the most important functions of DCs is to present antigens to lymphocytes to initiate adaptive immune responses (90–94). How DC death influences adaptive immunity during *Legionella* infection remains to be determined. Efferocytosis of dying DCs could provide antigens for uninfected bystander cells to take up and present to T cells as well as provide signals that direct lymphocyte differentiation. For example, phagocytosis of infected apoptotic cells by uninfected DCs induces a combination of cytokines that promotes T_H_17 cell differentiation (187). Engulfment of dying cells could also promote cross-priming of CD8^+^ T cells, as has been shown in other settings (188). Future work needs to be conducted to understand the consequences of DC death on infection outcome and its role in instruction of adaptive immunity.

Altogether, this study provides new insight into how DCs detect and restrict the intracellular bacterial pathogen *Legionella*. We find that early during infection, some DCs activate the caspase-11 and NLRP3 inflammasomes (Fig. 7). As the infection progresses, another group of DCs undergo effector-triggered apoptosis due to bacterial blockade of host protein synthesis, leading to decreased levels of the pro-survival proteins Mcl-1 and cFLIP (Fig. 7). Our findings lead us to propose a model where Bcl-2 family members and cFLIP act as guard proteins that respond to bacterial pathogen-mediated disruption of host translation in DCs (Fig. 7). These studies provide a foundation for understanding whether other bacterial pathogens similarly activate pyroptosis and effector-triggered apoptosis in DCs and other cell types.

## Materials and Methods

### Mice

All animal experiments were carried out in accordance with the Federal regulations set forth in the Animal Welfare Act (AWA), recommendations in the NIH Guide for the Care and Use of Laboratory Animals (189), and the guidelines of the University of Pennsylvania Institutional Animal Use and Care Committee. The animal protocol (#804928) was approved by the Institutional Animal Care and Use Committee (IACUC) at the University of Pennsylvania. All animals were housed and bred in specific-pathogen-free conditions. C57BL/6, *Nlrp3^−/−^* (190), *Casp1^−/−^* (191), and ASC-citrine reporter (192) mice were purchased from Jackson Laboratories. *Ripk3^−/−^* (193), *Ripk3^−/−^Caspase8^−/−^* (193, 194), *Gsdmd^−/−^* (191), *Gsdmd^−/−^Gsdme^−/−^* (47), *Casp1^−/−^ Casp11^−/−^* (58), and *Tnf^−/−^* (195) mice have been previously described and were bred in-house by the laboratory of Dr. Igor Brodsky.

### Mouse bone marrow-derived macrophage (BMDM) and dendritic cell (BMDC) culture

Bone marrow was harvested from mice and differentiated into macrophages by culture on non-tissue culture-treated (non-TC) dishes at 37°C in media comprising RPMI 1640, 30% L929 cell supernatant, 20% fetal bovine serum (FBS), 100 IU/mL penicillin, and 100 μg/mL streptomycin. Cells were fed at day 4 with 10mL of media. One day prior to infection (day 8), cells were plated in media comprising RPMI 1640, 15% L929 cell supernatant, and 10% FBS. Macrophages were plated at 1×10^5^ cells per well in 48-well TC treated plates and incubated at 37°C.

To generate BMDCs, bone marrow cells were differentiated by culture on non-tissue culture-treated dishes in media comprising RPMI 1640, 10% FBS, 50 µM 2-mercaptoethanol, 1% L-glutamine, 100 IU/mL penicillin, 100 μg/mL streptomycin, and 20 ng/mL recombinant GM-CSF (PeproTech 315-03). Cells were fed 5mL of media on days 3 and 5. One day prior to infection (day 8), cells were plated in media containing 10% FBS, 50 µM 2-mercaptoethanol, 1% L-glutamine and 5 ng/mL GM-CSF. Dendritic cells were plated at 2×10^6^ cells per well in 6-well non-TC plates, 2×10^5^ cells per well in 24-well non-TC plates, and 1×10^5^ cells per well in 48-well non-TC treated plates followed by incubation at 37°C.

### Bacterial culture

*Legionella pneumophila* strains derived from the Lp02 background (*thyA*; thymidine auxotroph) (64) were cultured on charcoal yeast extract agar (CYE) containing streptomycin for four days. Single colonies were isolated and cultured on CYE plates for 48 hours prior to use for infection in experiments. Lp02 Dot/Icm type IV secretion system-deficient (Δ*dotA*) and flagellin-deficient (Δ*flaA)* mutant strains were used and have been previously described (64, 196). The Lp02 Δ*7*Δ*flaA* strain lacked flagellin and 7 effectors that block host protein synthesis (Δ*lgt1* Δl*gt2* Δ*lgt3* Δ*sidI* Δ*sidL* Δ*LegK4* Δ*RavX*) (72). For experiments assessing bacterial replication, we used Lp02 strains carrying the plasmid pJB908, which encodes thymidine synthetase (18). Plasmid pJB908 containing Lgt2 or Lgt2* was transformed into competent Lp02 Δ*7*Δ*flaA* cells (18). Transformed cells were plated on CYE plates containing chloramphenicol. Colonies were screened via PCR to look for successful transformation of the plasmid. See Table S1 for a summary of the bacterial strains and plasmids used in this study along with their references.

### Bacterial infections

Heavy patches of *Legionella* grown on CYE agar plates for 48 hours were resuspended in sterile PBS. Cells were infected at a multiplicity of infection (MOI) of 50, unless otherwise stated. Infected cells were centrifuged and incubated at 37°C. For all experiments, control cells were mock-infected with PBS.

### Intracellular replication assays

Intracellular replication of *L. pneumophila* in BMDMs was measured as described previously (197) and modified slightly for BMDCs (96). DCs were infected at an MOI of 10 with *L. pneumophila*. Plates were centrifuged at 1200 rpm for 5 minutes and incubated at 37°C for 30 min. Cells were harvested from wells and anti-CD11c magnetic beads (Cat# 30-125-835; Miltenyi Biotec) were added. DCs were positively selected using MS columns (Miltenyi Biotec) in combination with a MiniMACS^TM^ magnetic separator, and extracellular bacteria were removed by washing the column with PBS containing 2% FBS. DCs were eluted from the column and 2×10^5^ cells were plated in non-tissue culture-treated 24-well plates. At indicated times after infection, adherent and non-adherent cells were harvested from wells and lysed with sterile H_2_O and combined with culture supernatants. Dilutions of the combined cell fractions and supernatants were plated on CYE plates to enumerate bacterial CFUs. Data are the mean CFUs recovered from three independent wells ± standard error of the mean (SEM). The fold increase in intracellular bacterial replication was calculated by determining the fold increase in CFUs at day 1 compared to the internalized bacterial CFUs at day 0.

### Cell death assays

Cells were infected as described above and assayed for cell death by measuring lactate dehydrogenase (LDH) activity in the supernatant using an LDH Cytotoxicity Detection Kit (Clontech) and normalized to mock-infected cells as well as cells treated with 1% Triton X-100 to establish maximum LDH release.

To measure cytotoxicity by uptake of propidium iodide (PI), cells were infected in 96-well black-walled tissue culture plates. At the time of infection, 5 μM PI was added to plate reader media (20 mM HEPES buffer and 10% FBS in Hank’s Balanced Salt Solution). Cells were then centrifuged at 1200 rpm for 5 minutes to infect and allowed to equilibrate to 37°C for 10 minutes. PI uptake into cells was then measured at an excitation wavelength of 530 nm and an emission wavelength of 617 nm. PI uptake was normalized to mock-infected cells and 1% Triton X-100-treated cells.

### Immunoblot analysis

Infected cells were lysed with 1x SDS/PAGE sample buffer. Protein samples were boiled for 5 min, separated by SDS/PAGE, and transferred to Immobilon-P membranes (Millipore). Samples were then probed with antibodies specific for Caspase-3 (Cell Signaling 9662), Caspase-7 (Cell Signaling 9492), Caspase-8 (Cell Signaling 4790), Caspase-9 (Cell Signaling 9508), BID (R&D MAB860), Mcl-1 (Cell Signaling 5453), Bcl-XL (Cell Signaling 2764), cFLIP (Cell Signaling 56343), Gasdermin-D (abcam 209845), Caspase-11 (Sigma C1354), and Caspase-1 (Genentech). As a loading control, all blots were probed with anti–β-actin (Cell Signaling 4967L). Detection was performed with HRP-conjugated anti-mouse IgG (Cell Signaling F00011), anti-rabbit IgG (Cell Signaling 7074S), or anti-rat IgG (Cell Signaling 7077).

Densitometry of pro-apoptotic proteins (Mcl-1, Bcl-XL, and cFLIP) was evaluated using ImageJ, normalizing to the corresponding β-actin band for that condition and timepoint, further normalized to the Mock infection.

### SUnSET assay

To assess global host protein synthesis, we utilized the previously described SUnSET assay (116). Briefly, BMDCs were infected at an MOI of 10 for 4 hours. Cells were treated with cycloheximide (1 μg/mL) as a positive control, which was added at the time of infection. After 4 hours of infection, puromycin was added to cells at 10 μg/mL for 1 hour. Lysates were harvested and processed as described above. Blots were probed for puromycin incorporation with an anti-puromycin antibody (DSHB Cat# PMY-2A4, RRID:AB_2619605). Area of staining was determined using ImageJ.

### ELISAs

Harvested supernatants from infected cells were assayed using ELISA kits for mouse IL-1α (R&D Systems) and IL-1β (BD Biosciences) following the manufacturer’s instructions.

### Fluorescence and confocal microscopy

BMDCs cells were seeded at a density of 8×10^5^ cells/mL on round, poly-l-ornithine-coated 12mm diameter glass coverslips (Electron Microscopy Sciences 72230-01) and allowed to adhere overnight in non-TC treated 24-well plates. Cells were then transfected with *Legionella* strains at an MOI of 50. At the indicated timepoints, cells were washed 3 times with PBS and fixed with 4% paraformaldehyde. Cells were stained with primary antibody against rabbit anti-cleaved caspase-3 (Cell Signaling 9664) followed by anti-rabbit AF647 (Invitrogen A21245). Following nuclear counterstain with Hoechst 33342 (1 μg/mL; Thermo Fisher #62249), cells were mounted on glass slides with ProLong Glass (Invitrogen P36882) and dried overnight. Slides were imaged using a Zeiss LSM 980 confocal microscope at a single z-plane per field with lasers optimized for Cy5 (far-red), citrine (yellow), and CellTracker Biolet (Blue) emission spectra, through a ×63 objective. The presence of ASC specks and cleaved caspase-3 was assessed in individual cells. A speck was defined as a distinct high-fluorescent perinuclear cluster of citrine signal. Each experiment was analyzed for an average of 100-150 cells. Three independent experiments were quantified and graphed.

### Statistical analyses

Graphing and statistical analysis were carried out in GraphPad Prism 10. In comparisons between two groups, unpaired Student’s t-test was utilized to determine significance. In comparisons between more than two groups, two-way ANOVA was utilized to determine significance, with Tukey HSD test following up. Difference considered significant when the P value is < 0.05.

## Acknowledgements

We thank members of the Shin and Brodsky laboratories for helpful scientific discussions. We also thank Drs. Tzvi Pollock, Igor Brodsky, and Heather L. Rossi for critical reading of the manuscript. We thank Mark A. Boyer for his help with the breeding of C57BL/6 mice and Beatrice Hermann and Ronit Schwartz from the Brodsky lab for providing *Gsdmd^−/−^*, *Gsdmd^−/−^Gsdme^−/−^*, and *Casp1^−/−^Casp11^−/−^* mice. We thank the Vance laboratory for providing *Legionella* strains. We would also like to thank the Cell & Developmental Biology Microscopy Core (RRID SCR_022373) at the Perelman School of Medicine at the University of Pennsylvania for access and usage of the Zeiss LSM 980 microscope. We thank the University of Pennsylvania Summer Undergraduate Internship Program (SUIP) for supporting Madison Dresler and Karly Kammann, the University of Pennsylvania Post-Baccalaureate Research Education Program for supporting Frankie D. Boyer, and the University of Pennsylvania Postdoctoral Opportunities in Research and Teaching for supporting Dr. Kimmie A. Wodzanowski. This work is supported in part by National Institutes of Health grants R01AI118861 (S.S.), R21AI12178 (S.S.), K12GM081259 (K.A.W.), NIH T32 AI141393-03 (J.Z), R25GM071745-19 (F.D.B.), and R25HL084665 (M.V.D.), National Science Foundation Graduate Fellowship DGE-1845298 (V.R.V.M), American Heart Association Predoctoral Fellowship (J.D.), and a Burroughs-Welcome Fund Investigators in the Pathogenesis of Infectious Diseases.

---

**Figure S1. Caspase-8 is not the sole driver of apoptosis in *Legionella*-infected dendritic cells.** WT, *Ripk3^−/−^*, or *Ripk3^−/−^Caspase8^−/−^* BMDCs were mock-infected or infected with T4SS^+^ *Legionella* at a MOI of 50 for 4hr. Immunoblot analysis was performed on cell lysates for caspase-3, caspase-7, and caspase-8 with β-actin as a loading control. For cleaved products, lanes from one membrane have been cropped and moved to depict the appropriate conditions. No changes were made to the original image during the editing. long exp., long exposure. Data shown are representative of at least two independent experiments.

**Figure S2. Effector-triggered apoptosis does not account for all cell death occurring in dendritic cells.** BMDCs were infected with T4SS^+^ or T4SS^+^Δ*7 Legionella* at an MOI of 50 and cytotoxicity was measured using PI uptake assay. Data shown are representative of at least three independent experiments.

**Figure S3. *Legionella*-mediated block in host protein synthesis decreases IL-1 secretion in dendritic cells.** WT BMDCs were mock-infected or infected with T4SS^+^ or T4SS^+^Δ*7 Legionella* at MOI 50. Cytokine release was measured at 4hr using ELISA. Data shown are representative of at least three independent experiments.

**Figure S4. TNF promotes pyroptosis but not cell-extrinsic apoptosis in dendritic cells during *Legionella* infection.** (A-B) WT or *Tnf^−/−^* BMDCs were infected with T4SS^+^ *Legionella* at an MOI of 50. Cytotoxicity was measured by PI uptake assay (A) or LDH release assay at 4hr (B). (C) WT or *Tnf^−/−^* BMDCs were mock-infected (represented as “-“) or infected with T4SS^+^ *Legionella* at an MOI of 50 for 1, 2, 3, or 4 hr post-infection. Immunoblot analysis was performed on cell lysates for GSDMD, caspase-11 and β-actin as a loading control. (D) WT or *Tnf^−/−^* BMDCs were mock-infected (represented as “-“) or infected with T4SS^+^ *Legionella* at an MOI of 50 for 4 hr post-infection. Immunoblot analysis was performed on cell lysates for caspase-8, caspase-3, caspase-7, and β-actin as a loading control. For cleaved products, lanes from one membrane have been cropped and moved to depict the appropriate conditions. No changes were made to the original image during the editing. long exp., long exposure. hpi, hours post-infection. Data shown are representative of at least two (C,D) or three (A,B) independent experiments. Graphs show the mean ± SEM of triplicate wells. Data were analyzed by two-way ANOVA with Sidak’s multiple comparisons test; ****, P < 0.0001.

**Table S1. Summary of the bacterial strains and plasmids used in this study and their references.**

