## Supplemental figures for "Dendritic cells activate pyroptosis and effector-triggered apoptosis to restrict *Legionella* infection"

Figure S1

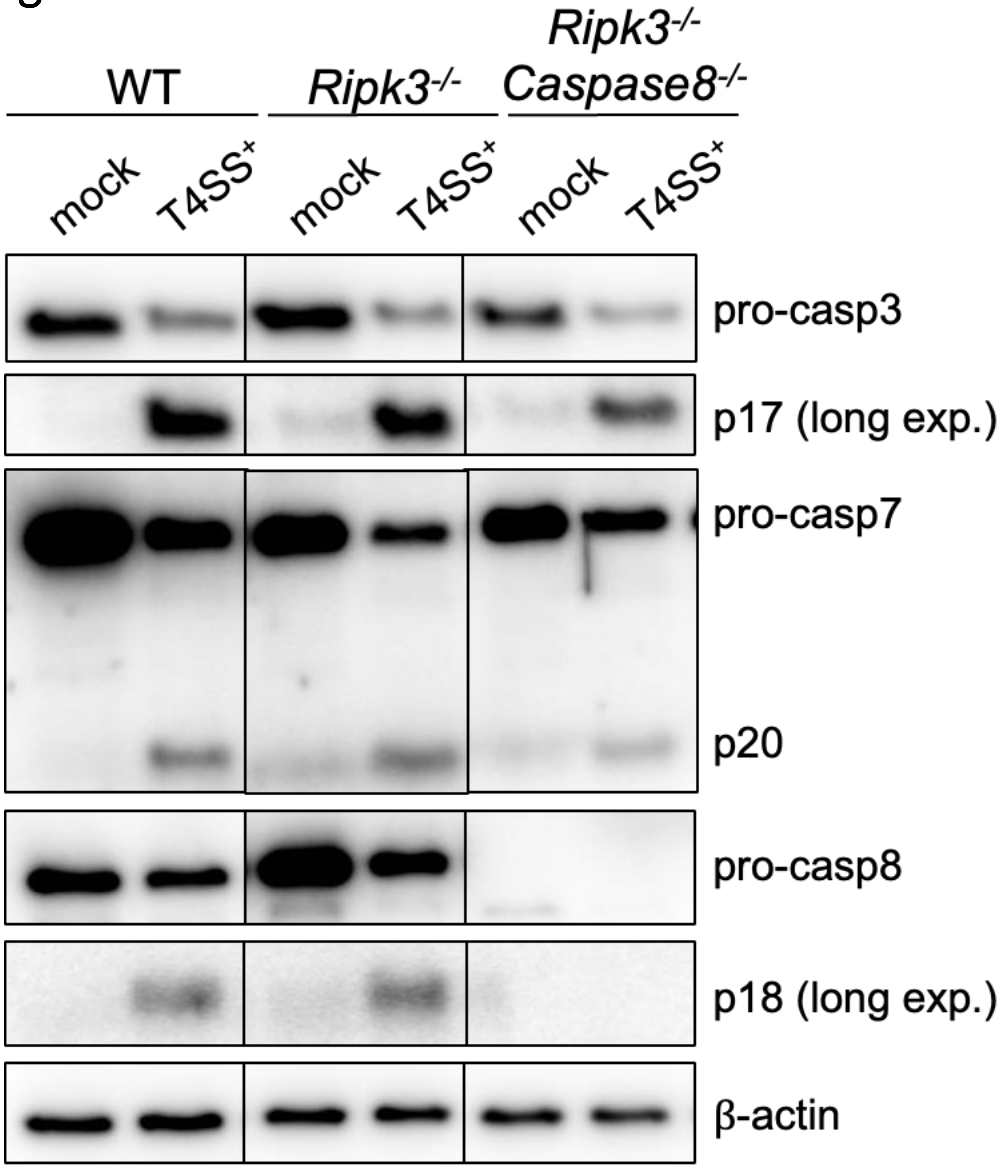
WT
Ripk3-/-
Ripk3-/- Caspase8-/-
mock
T4SS+
mock
T4SS+
mock
T4SS+
pro-casp3
p17 (long exp.)
pro-casp7
p20
pro-casp8
p18 (long exp.)
β-actin

Figure S2

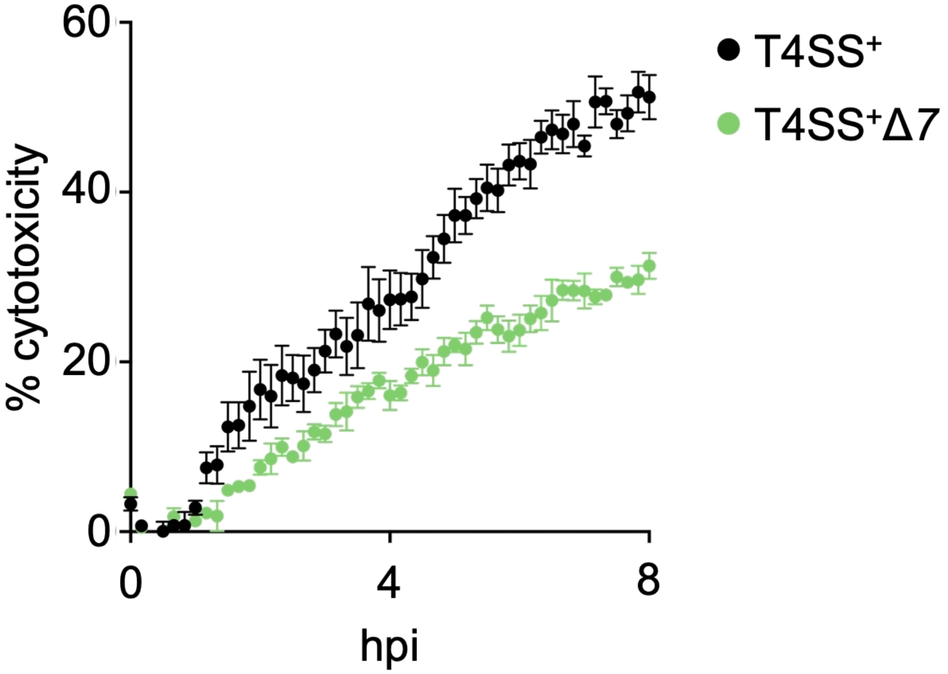

Figure S3

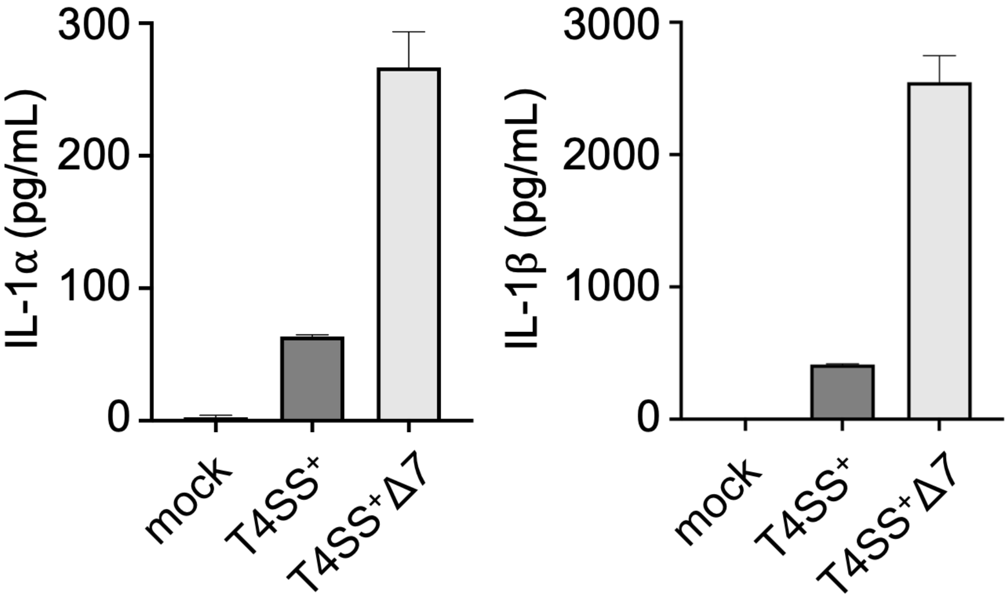

IL-1α (pg/mL)
300
200
100
0
mock
T4SS⁺
T4SS⁺Δ7
IL-1β (pg/mL)
3000
2000
1000
0
mock
T4SS⁺
T4SS⁺Δ7

Figure S4

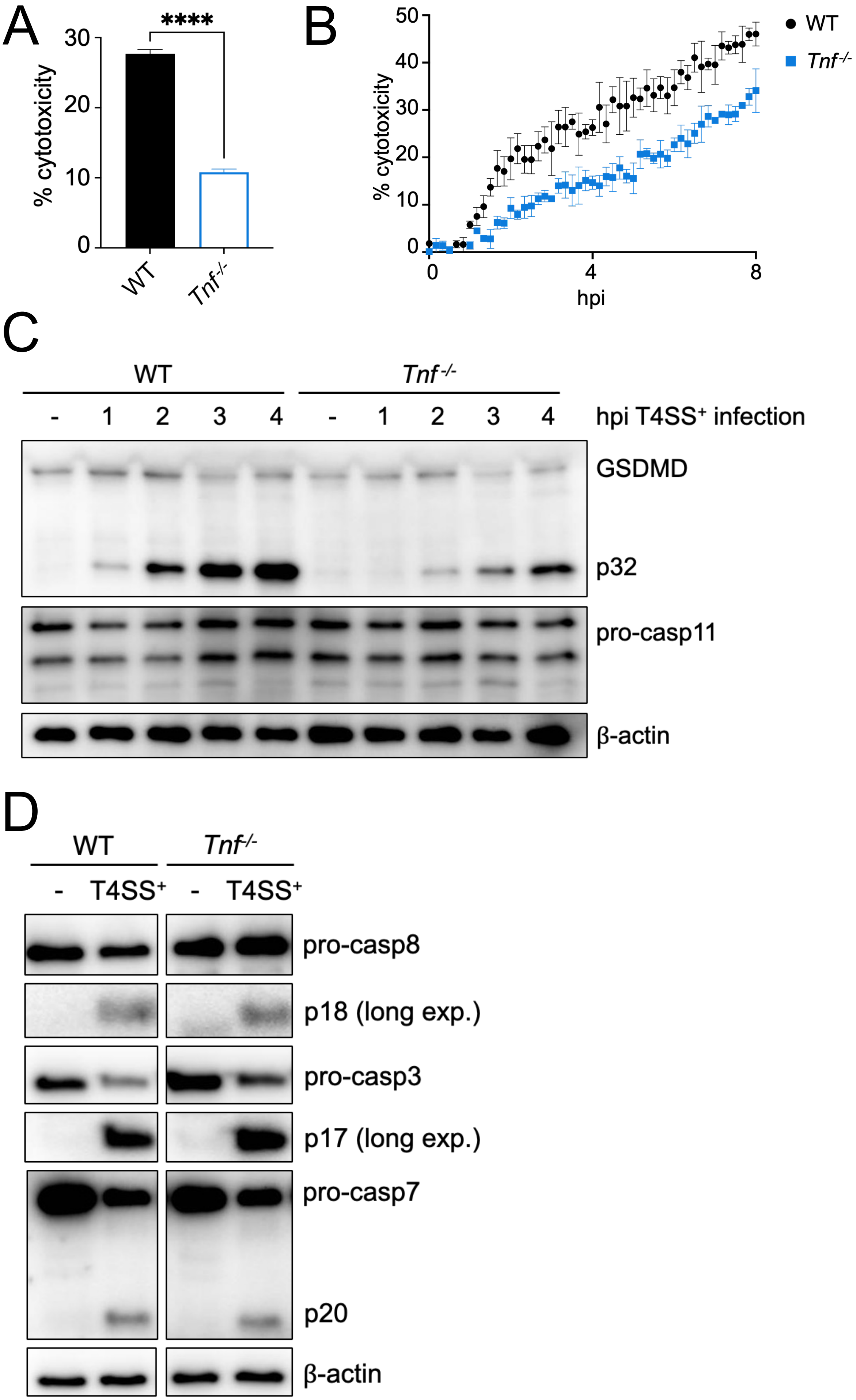
A
****
% cytotoxicity
0
10
20
30
WT
Tnf-/-
B
% cytotoxicity
0
10
20
30
40
50
hpi
0
4
8
WT
Tnf-/-
C
WT
Tnf-/-
- 1 2 3 4 - 1 2 3 4
hpi T4SS+ infection
GSDMD
p32
pro-casp11
β-actin
D
WT
Tnf-/-
- T4SS+ - T4SS+
pro-casp8
p18 (long exp.)
pro-casp3
p17 (long exp.)
pro-casp7
p20
β-actin

Table S1

| Bacterial strain or plasmid | References |
|---|---|
| *Legionella pneumophila* Lp02 Δ*flaA* | Berger KH, Isberg RR. 1993<br>Ren T, Zamboni DS, Roy CR, Dietrich WF, Vance RE. 2006 |
| *Legionella pneumophila* Lp02 Δ*dotA*Δ*flaA* | Berger KH, Isberg RR. 1993<br>Ren T, Zamboni DS, Roy CR, Dietrich WF, Vance RE. 200 |
| *Legionella pneumophila* Lp02 Δ*7*Δ*flaA* | Barry KC, Fontana MF, Portman JL, Dugan AS, Vance RE. 2013. |
| pJB908 | Fontana MF, Banga S, Barry KC, Shen X, Tan Y, Luo Z-Q, Vance RE. 2011. |
| pJB908 containing Lgt2 | Fontana MF, Banga S, Barry KC, Shen X, Tan Y, Luo Z-Q, Vance RE. 2011. |
| pJB908 containing catalytically inactive Lgt2 | Fontana MF, Banga S, Barry KC, Shen X, Tan Y, Luo Z-Q, Vance RE. 2011. |
